# Type IV pilin post-translational modifications modulate materials properties of bacterial colonies

**DOI:** 10.1101/389882

**Authors:** R. Zöllner, T. Cronenberg, N. Kouzel, A. Welker, M. Koomey, B. Maier

**Author notes:** equal contribution (R. Zöllner, T. Cronenberg).

## Abstract

Bacterial type 4 pili (T4P) are extracellular polymers that initiate the formation of microcolonies and biofilms. T4P continuously elongate and retract. These pilus dynamics crucially affects the local order, shape, and fluidity of microcolonies. The major pilin subunit of the T4P bears multiple post-translational modifications. By interfering with different steps of the pilin glycosylation and phosphoform modification pathways, we investigated the effect of pilin post-translational modification on the shape and dynamics of microcolonies formed by *Neisseria gonorrhoeae*. Deleting the phosphotransferase responsible for phosphoethanolamine modification at residue serine 68 (S^68^) inhibits shape relaxations of microcolonies after pertubation and causes bacteria carrying the phosphoform modification to segregate to the surface of mixed colonies. We relate these mesoscopic phenotypes to increased attractive forces generated by T4P between cells. Moreover, by deleting genes responsible for the pilin glycan structure, we show that the number of saccharides attached at residue serine 63 (S^63^) affect the ratio between surface tension and viscosity and cause sorting between bacteria carrying different pilin glycoforms. We conclude that different pilin post-translational modifications moderately affect the attractive forces between bacteria but have severe effects on the materials properties of microcolonies.

## Introduction

Many if not most bacteria live in structured communities called biofilms. The first stage of biofilm formation is the aggregation of bacteria into small clusters or (micro)colonies. Interestingly, the size and shape of colonies can respond to changes in the environment. For example, *Staphylococcus aureus* forms large biofilm structures in response to antibiotic treatment (1). *Pseudomonas aeruginosa* enlarges the size of colonies in the presence of predators (2). Moreover, bacteria tend to increase the sufarce area of biofilms in response to nutrient limitation (3, 4) or oxygen depletion (5). Thus it is well established that bacteria control biofilm architecture. On the other hand, very little is known about how bacteria remodel colonies and biofilms.

Many bacterial species employ extracellular proetinaceous appendages called type 4 pili (T4P) for initiating cellular aggregation (6-8). T4P are dynamic polymers that extend by polymerization and retract by depolymerization (9) (10). The main constituent of T4P is the major pilin (encoded by *pilE* in the organism studied here). Elongation and retraction are driven by dedicated ATPases (11) (12). During the process of retraction, the T4P is capable of generating very high force (13) (14). Interestingly, the fact that the T4P elongate and retract dynamically affects the shape, dynamics, and sorting behavior of bacterial colonies (7, 15, 16). *Neisseria gonorrhoeae* and *Neisseria meningiditis* are very good systems for studying the role of T4P in colony formation and dynamics. Non-piliated *N. gonorrhoeae* do not aggregate, indicating that the T4P is the major factor controlling the interactions between bacteria in colonies (16). T4P retraction has been shown to affect colony shape. *N. gonorrhoeae* usually form spherical colonies. However, the shapes of mutants colonies generating T4P that are incapable of retraction are undefined (7). By analogy to liquids, bacteria generating attractive force produce surface tension the causes colonies to assume a spherical shape (15, 17). Moreover, bacteria show local liquid-like order (17). When colonies are deformed by fusion of two colonies, their shape relaxes towards a sphere with higher radius. The dynamics of shape relaxation is liquid-like for wild type gonococci (17). For colonies formed by bacteria with non-retractile T4P, no shape relaxation was observed and the single cell dynamics within the colony was severely reduced (15). Fluidization of colonies in association with T4P retraction has been suggested to be important for colonization of blood capillary network during infection (15). T4P-mediated dynamics also enables sorting of cells with respect to differential densities of T4P and interaction forces (16). When bacteria with different T4P densities are mixed, or when bacteria lose T4P as a consequence of pilin antigenic variation, they sort to the front of a growing colony (18).

Many T4P major pilins can be post-translationally modified. For *N. gonorrhoeae*, the pilin is *O*-glycosylated at S^63^ (19) and carries a phosphoform-modification at S^68^ (20). In particular, wildtype pilins minimally bear a di-N-acetylbacillosamine sugar (diNAcBac) generated by (21) generated by *O*-linked protein glycosylation (*pgl*) pathway (Fig. 1a). Biosynthesis of the pilin glycan involves a lipid-linked monosaccharide precurser. The *pglA* gene encodes for a glycosyltransferase that adds a galactose onto the basal diNAcBac (21). PglE acts as a glycosyltransferase that adds the second galactose to make diNAcBac-Gal-Gal trisaccharide (21). Next, the flippase PglF is involved in membrane translocation or flipping of the lipid-attached carbohydrate. Finally, the oligosaccharyltransferase PglO carries out *en bloc* glycan transfer to the pilin (21). For different species or isolates, the pilin can bear different oligosaccharide modifications (22). For example, *pglA* and *pglE* are phase-variable (21). Due to a homopolymeric stretch in its DNA sequence, the gene undergoes frameshift mutations at a high rate, switching the gene between the ON- and the OFF-state. The *N. gonorrhoeae* pilin used in this study additionally carries a phosphoethanolamine moiety at S^68^ (20). The phosphotransferase PptA is required for this post-translational modification (23). There is evidence, that both glyco- and phosphophorm modifications affect colony dynamics. In a related system, *N. meningiditis* was reported to upregulate expression of the phosphotransferase gene responsible for pilin phosphoform modification during infection of host cells. This event was associated with colony dispersal (24). In line with these obervation, computer simulations suggest that the post-translational modifications affect the physical interactions between T4P in pilus bundles (24). Recently, we showed that deletion of the flippase *pglF* altered the T4P-mediated rupture forces between gonococci (16). As predicted by the differential force of adhesion hypothesis (25), bacteria with glycosylated T4P segregated from bacteria where the glycosylation pathway was blocked (16). These studies suggest that post-translational modification of pilin strongly affects colony structure and dynamics.

**Fig. 1.**
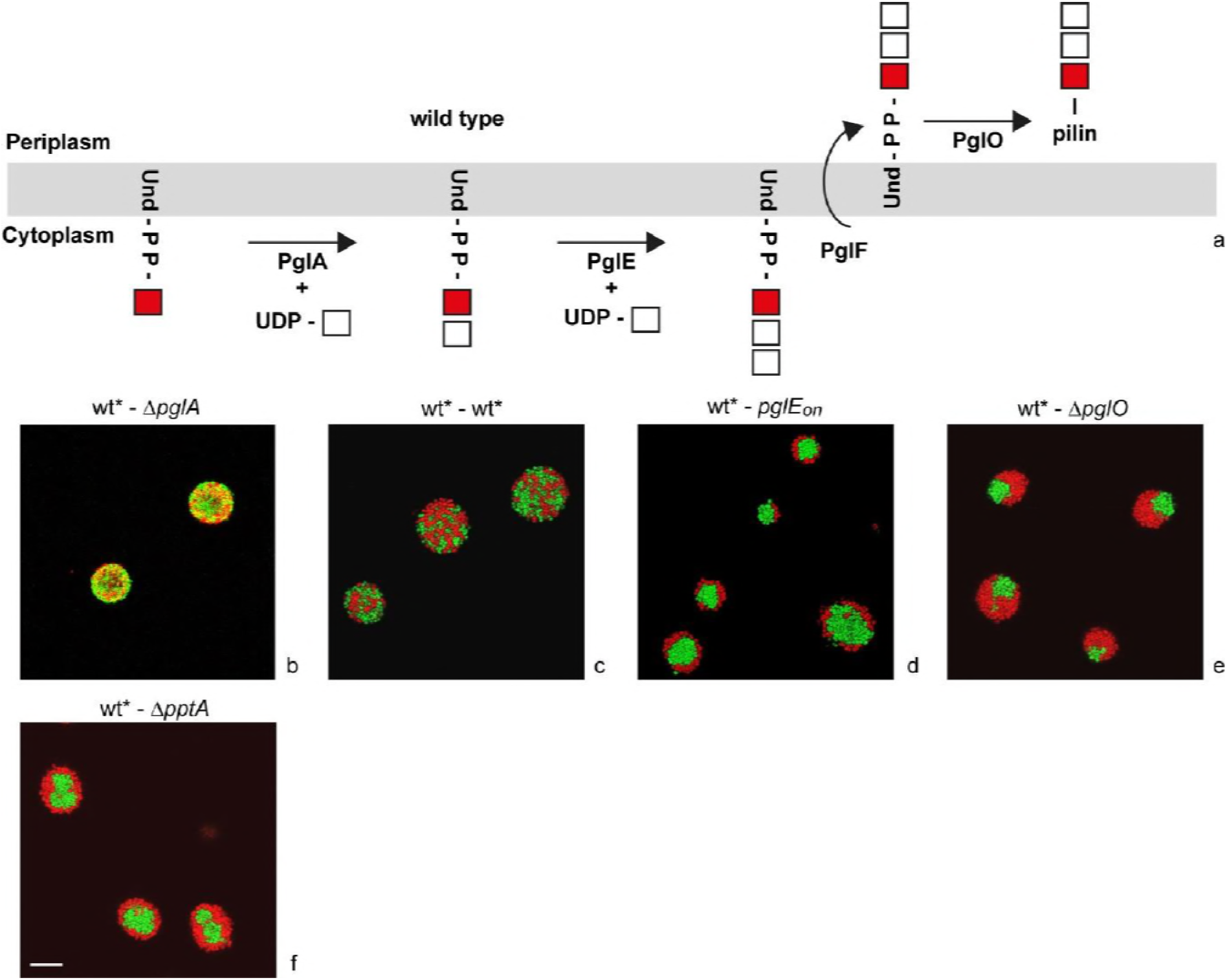
Different pilin post-translational modifications trigger cell sorting in gonococcal microcolonies. a) Current model of pilin glycosylation pathway (adapted from (21)). *mcherry*- expressing wt* (Ng170) cells were mixed with *gfp*-expressing b) *∆pglA* (Ng141), c) wt* (Ng151), d) *pglE_on_* (Ng157), e) *∆pglO* (Ng140), and f) *∆pptA* (Ng142), respectively, at a 1 : 1 ratio. After 5 h of incubation, colonies were imaged with confocal microscopy. In all strains, the G4-motive was deleted. Scale bar 10 μm.

In this study, we investigate the influence of different types of pilin post-translational modifications on attractive forces between bacteria, cellular sorting, and the dynamics of colony remodeling. Inhibiting or activating different steps of the pilin glycosylation pathway causes segregation of the respective mutants from the wild type bacteria in mixed colonies and affect the shape relaxation dynamics of microcolonies after perturbation. Inactivation of the gene whose product is responsible for phosphophorm modification causes very robust sorting behavior and inhibits shape relaxations, suggesting that enhanced T4P-T4P interactions increase the viscosity inside colonies. Our study relates changes in molecular interactions between T4P caused by different pilin post-translational modifications to the mesoscopic spatio-temporal dynamics of bacterial colonies.

## Materials and Methods

### Growth conditions

Gonococcal base agar was made from 10 g/l BactoTM agar (BD Biosciences, Bedford, MA, USA), 5 g/l NaCl (Roth, Darmstadt, Germany), 4 g/l K2HPO4 (Roth), 1 g/l KH2PO4 (Roth), 15 g/l BactoTM Proteose Peptone No. 3 (BD), 0.5 g/l soluble starch (Sigma-Aldrich, St. Louis, MO, USA)) and supplemented with 1% IsoVitaleX: 1 g/l D-Glucose (Roth), 0.1 g/l L-glutamine (Roth), 0.289 g/l L-cysteine-HCL×H20 (Roth), 1 mg/l thiamine pyrophosphate (Sigma-Aldrich), 0.2 mg/l Fe(NO3)3 (Sigma-Aldrich), 0.03 mg/l thiamine HCl (Roth), 0.13 mg/l 4-aminobenzoic acid (Sigma-Aldrich), 2.5 mg/l β-nicotinamide adenine dinucleotide (Roth) and 0.1 mg/l vitamin B12 (Sigma-Aldrich). GC medium is identical to the base agar composition, but lacks agar and starch.

### Bacterial biofilm growth conditions

Each biofilm was grown within ibidi μ-Slides I^0.8 Luer flow chambers for 5 h at constant nutrient flow of 3 ml/h by using a peristaltic pump (model 205U; Watson Marlow, Falmouth, United Kingdom). Bacteria from overnight plates of each strain were resuspended in GC medium to an optical density at 600nm (OD600) of 0.1 and 300 μl of cell suspension was inoculated into the flow chambers. The bacteria were left for 1 h at 37°C to allow for attachment to the glass surface. After 1 h, the flow was switched on.

### Bacterial strains

All bacterial strains were derived from the gonococcal strain MS11 (VD300) (Table 1). In all strains, we deleted the G4-motif upstream of the pilin gene by replacing it with the *aac* gene (conferring resistance against apramycin). The G4-motif is essential for antigenic variation of the major pilin subunit (26). Pilin antigenic variation modifies the primary sequence of the pilin gene. Since the composition of amino acids can affect the pilus density and potentially the rupture force between pili, antigenic variation is likely to generate heterogeneity and distract form the major topic of this study. Since there is no reason to expect that deletion of the G4 motif affects the structure of the biofilm, the strain carrying the G4 deletion is labeled wt* throughout the manuscript.

**Table 1.** Strains used in this study

| Strain | Relevant genotype | Source/Reference |
| --- | --- | --- |
| *wt* VD300 (Ng002) | wild type, *opa*- selected | (16) |
| *wt* * (Ng150) | *G4::aac* | (18) |
| *wt* * (Ng151) | *iga::P<sub>pilE</sub> gfpmut3 ermC*<br>*G4::aac* | (17) |
| *wt* * (Ng170) | *lctp:mcherry aadA:aspC*<br>*G4::aac* | (18) |
| Δ*pglA* (Ng141) | *pglA::kan*<br>*iga::P<sub>pilE</sub> gfpmut3 ermC*<br>*G4::aac* | This study, (21) |
| *pglE<sub>on</sub>* (Ng157) | *iga::P<sub>pilE</sub> gfpmut3 ermC*<br>*G4::aac* | This study |
| Δ*pglO* (Ng140) | *pglO::kan*<br>*iga::P<sub>pilE</sub> gfpmut3 ermC*<br>*G4::aac* | This study, (21) |
| Δ*pptA* (Ng142) | *pptA::kan*<br>*iga::P<sub>pilE</sub> gfpmut3 ermC*<br>*G4::aac* | This study, (23) |

Strains *∆pglA* (Ng141), *∆pglO* (Ng140), and *∆pptA* (Ng142) were generated by transforming strain Ng151 with chromosomal DNA of strains KS141 (21), KS145 (21), and KS9 (23), respectively, and selecting for kanamycin resistance. *pglE* carries 5 CAAACAC copies and is therefore prone to phase variation. As consequence, *pglE* is in its OFF-stain in strain Ng151. Thus the latter was transformed with a PCR product generating strain *pglE_on_* (Ng157) where *pglE* is in its ON-state. Fragment 1 (upstream of *pglE*) was amplified using the primers NK167 atgccgtctgaaCAGTGAAAATCCCGAAGACATCAAAGC and NK169 ccccagctggcaattccggTTATTTGCCCGCTTTGAGCCCTTGC. Fragment 2 contained the kanamycin resistance cassetts and was amplified from pGCC6 (27) using NK170 GCAAGGGCTCAAAGCGGGCAAATAAccggaattgccagctgggg and NK172 CAGACGGCAGGGATTCCGGtcagaagaactcgtcaagaaggcgatag. Fragment 3 contained the functional *pglE* gene and was amplified from KS142 (21) where *pglE* was in the ON-state using NK171 ctatcgccttcttgacgagttcttctgaCCGGAATCCCTGCCGTCTG and NK173 ttcagacggcatTTAAATATTCCCCCTGATTGCTTTTAAAATCCTG. The three segments were fused by PCR. Correct insertion was verified by PCR and sequencing (GATC, Konstanz).

### Confocal microscopy

After 5h of growth, biofilms were fixed with 4% Formaldehyde in 1x PBS for 15 min at RT and washed with GC medium. Colony images were acquired using a Leica TCS SP8 confocal laser scanning microscope (CECAD Imaging Facility) with a 63x, 1.4 NA, oil immersion objective lens. The excitation wave lengths were 488 nm and 587 nm. The microcolony in Fig. 1b was imaged with a Nikon TE2000 C1 microscope.

### Measurement of bacterial interaction forces

In order to characterize single cell interactions, we followed a previously developed protocol (17). The optical trap was installed in an inverted microscope (Nikon TE2000 C1) (28). The trapping laser (20I-BL-106C, Spectra Physics, 1064nm, 5.4W) was directed into a water-immersion objective (Nikon Plan Apochromate VC 60x N.A. 1.20). Manipulation of the trap position was done with a two-axis acousto-optical deflector (DTD-274HD6 Collinear Deflector, IntraAction Corp., USA). Bacterial interaction was recorded with a CCD camera (sensicam qe, PCO, Kelheim, Germany). The optical trap was calibrated via the equipartition and drag force methods. The average stiffness is 0.11 (± 0.01) pN/nm. The linear regime extends up to 80pN.

### Analysis of colony fusion dynamics

Colony fusion dynamics were analyzed as described in Welker et al. (17). A custom build flow chamber was mounted into an inverted microscope (Nikon TI). The chamber consisted out of two cover slips and a holder where the bottom coverslip was coated with BSA to prevent strong adhesion of the bacteria. Bacteria were resuspended in GC containing 1% Isovitalex, diluted to an OD of 0.25 and injected into the chamber. Images were analyzed using Matlab and the DIPimage toolbox. For the analysis, only colonies with a volume v < 10^4^ μm^3^ were taken into account. The contour of the object was extracted and fitted by an ellipse for every single time point to extract the ratio *f* = *b/a* between minor axis *b* and major axis *a*. We used a fluid model (29, 30) to extract the ratio between surface tension σ and viscosity *η* out of the shape relaxation from an ellipsoid towards a sphere. The rate of rounding up is

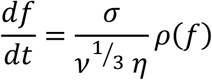

where *v =* 4/3 *nab^2^*, and

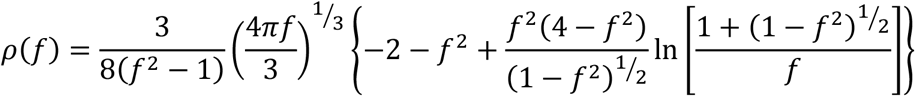

The expression for the time it takes for an ellipse to relax from f0 to f_1_

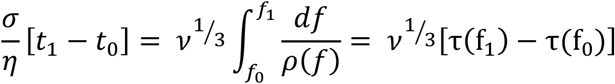

was integrated numerically with f_0_ = 0.71. The lower limit f0 was chosen such that the aggregates were in good agreement with the shape of an ellipse. The ratio between surface tension *σ* and viscosity *η* was calculated by a linear fit.

## Results

### Differential post-translational modification of T4P impacts cell sorting

Previously, we showed that gonococci sorted with respect to the number of T4P (16). Furthermore, inhibition of *O*-linked pilin glycosylation triggered sorting. Here, we investigate whether different post-translational pilin modifications induce cell sorting. To this end, we generated *gfp*-expressing gonococcal strains with inactivating deletions in genes that encode for enzymes involved in pilin post-translational modifications (Fig. 1a). These strains were mixed at a 1 : 1 ratio with wild type (wt*) cells, incubated for 5 h, and subsequently imaged using confocal microscopy (Fig. 1b). In *N. gonorrhoeae*, the gene encoding for the major pilin, *pilE*, undergoes antigenic variation (26). As a consequence, the primary structure of *pilE* varies. The G4 motif upstream of the *pilE* openreading frame is essential for pilin antigenic variation (26). To avoid potential effects on T4P-T4P interactions caused by pilin antigenic variation, in all strains used in this study the G4 motif was deleted.

The wildtype pilin in the parental background bears a disaccaride composed of a galactose residue linked to diNAcBac forming diNAcBac-Gal (21). When *gfp-* and mcherry-expressing wt* cells were mixed, there was no evidence for cell sorting as expected (Fig. 1c). Next, we mixed a *gfp-* expressing *∆pglA* strain with the mcherry-expressing wt* strain (Fig. 1b). The PglA glycosyltransferase adds the galactose onto the basal diNAcBac (21). Thus deletion of *pglA* results in a pilin which is modified by a truncated diNAcBac monosaccharide. We found no evidence for segregation of the *∆pglA* strain from the wt* strain (Fig. 1b), indicating that gonococci with mono- and disaccharide pilin modifications mix well. Next, we investigated the effect of pilins bearing a trisaccharide. PglE acts as a glycosyltransferase that adds the second galactose to make the diNAcBac-Gal-Gal trisaccharide (21). The *pglE* gene is phase-variable and in our wt* strain, the gene is in its OFF-state. We engineered a stable *pglE_on_* strain where *pglE* is in its ON-state. Strain *pglE_on_* segregated to the center of the mixed colony (Fig. 1d). Thereby wt* cells spread fairly homogeneously on the interface to the core of the colony formed by *pglE_on_* cells. Finally, we confirmed that deletion of *pglO* caused sorting (Fig. 1e). PglO is the enzyme responsible for *en bloc* glycan transfer to the pilin (21). Thus, deletion of *pglO* results in non-glycosylated pilin. We found that *∆pglO* mutants and wt* cells sorted. Interestingly, wt* cells partially spread around the core formed by *∆pglO* cells. This behavior is reminiscent of *∆pglF* deletion mutations reported previously (16).

In addition to an *O*-linked saccharide at S^63^ the strain used here carries an additional post-translational modification of phosphoethanolamine at S^68^ (20). The pilin phosphoform transferase A (PptA) is required for the phosphoform-modification (23). To address the question as to whether the latter pilin modification caused segregation within colonies, we mixed wt* cells with *∆pptA* cells. wt* cells sorted to the surface of the mixed colony and spread fully over the *∆pptA* core (Fig. 1f).

We conclude that sorting with respect to differential pilin post-translational modifications is a general phenomenon in *N. gonorrhoeae*.

### T4P post-translational modification reduces the strength of cell-cell attraction and affects the binding probability

We intended to link the sorting behavior at the colony level to T4P-mediated interaction forces between bacteria. To this end, we used a double laser trap for measuring the rupture forces between T4P emanating from different cells (Fig. 2a) (17). Here, *mcherry-expressing* wt* cells were mixed with g¿p-expressing mutant cells. We trapped a monococcus within each laser focus. The laser traps had a distance of 2.84 μm. When T4P from different cells bind to each other and at least one of the pili retracts, the cell bodies attract each other. This attraction is detected as a deflection *d* from the centers of the laser traps. Each event starts with a retraction. Eventually, the bond between T4P ruptures and the cell bodies move back towards to centers of the traps. The pair of cells was always in one of the following states: detached, retracting, elongating, or pausing. Each pair of cells was oberserved for ~ 1min. Subsequently, fluorescence microscopy was used to find out which pair of cells (wt* - wt*, wt* - mutant, or mutant - mutant) have been characterized.

**Fig. 2.**
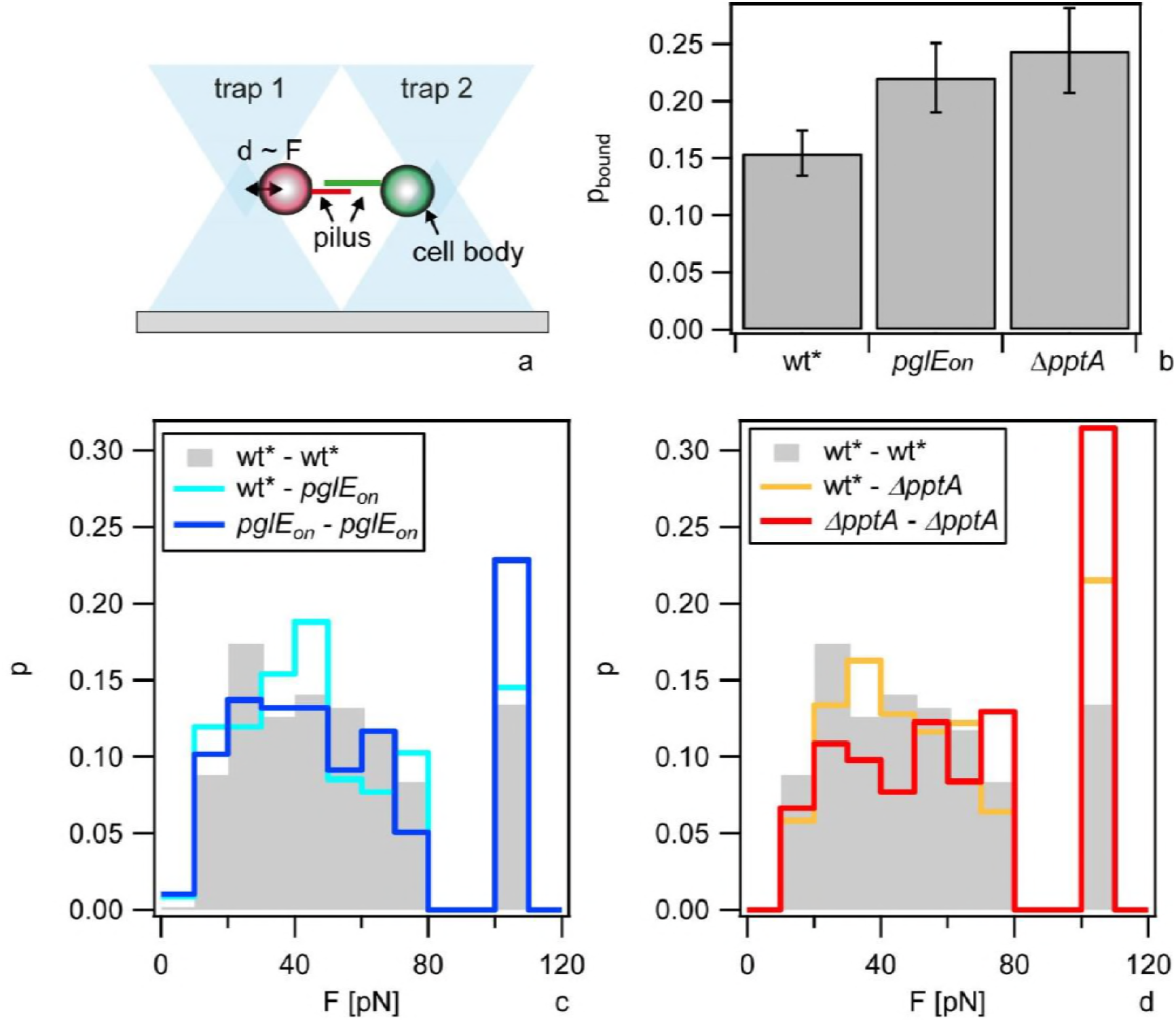
A dual laser trap was used to determine the rupture force between T4P of two cells. a) Sketch of the setup. mcherry-expressing wt* cells and *gfp*-expressing mutants cells were mixed. In each laser trap, a single cell was immobilized. Deflection *d* (~ force *F*) of both cells from the centers of the traps was recorded as a function of time. After recording was finished, fluorescence microscopy was used to determine which cell types were trapped. b) Probability that at least one pair of T4P from different cells are attached to each other. Error bars: bootstrapping. c) Rupture forces between grey: wt*(Ng 170) and wt* (Ng170), cyan: wt* (Ng170) and *pglE_on_* (Ng157), blue: *pglE_on_* (Ng157) and *pglE_on_* (Ng157). The rupture forces of *pglE_on_* /*pglE_on_* are larger than wt*/wt* at a significance level of P = 0.1 (KS-test). d) Rupture forces between grey: wt* (Ng170) and wt* (Ng170), orange: wt* (Ng170) and *∆pptA* (Ng142), red: *∆pptA* (Ng142) and *∆pptA* (Ng142). The rupture forces of *∆pptA* / *∆pptA* are larger than wt*/wt* at a significance level of P = 10^-8^ (KS-test). All data for forces exceeding 80 pN were binned into a single bin at 100 pN because the force increased in a non-linear fashion as a function of the deflection.

The most interesting quantities for linking T4P dynamics to cell sorting and colony behavior are the probability *p_bound_* that two cells are bound to each other via T4P and the rupture force between pili. *p_bound_* is different from the binding rate; it measures the probability that two cells trapped at a distance of 2.84 μm are bound to each other via their T4P at any point of time. We found that the probability of wt* - wt* binding was p_bound wt*-wt*_ = 0.15 ± 0.02 (Fig. b). The probabilities for*pglE_on_* -*pglE_on_* and *∆pptA - ∆pptA* binding were significantly higher with p_bound *pglE_on_-pglE_on_*_ = 0.22 ± 0.03 and p_bound *∆pptA-∆pptA*_ = 0.24 ± 0.04. The rupture forces were broadly distributed. Since the force of the laser trap decreases linearly with the deflection of the cell bodies only for forces below F_rupture_ = 80 pN, all data exceeding this value were binned into a single bin at 100 pN. The rupture forces between *pglE_on_* - *pglE_on_* were shifted to slightly higher values compared to the wt* - wt* rupture forces (Fig. 2c). However, the significance of this shift is not very high. By contrast, *∆pptA - ∆pptA* rupture forces exceeded the wt* - wt* rupture forces considerably (Fig. 2d). The distribution of *∆pptA -* wt* rupture forces was situated between the distributions for *∆pptA - ∆pptA* and wt* - wt*. Deletion of*pglA* did not cause cell sorting and was not investigated further. Previously, we showed that inhibition of pilin glycosylation affected the rupture force between T4P (16). Therefore, we did not repeat the experiment with the *∆pglO* strain here.

In summary, both the addition of a third hexose to the saccharide at S^63^ and inhibition of pilin phosphoform-modification at S^68^ increase the T4P-mediated probability that gonococci are bound. In addition, inhibition of pilin phosphoform-modification increases the rupture force between T4P.

### T4P post-translational modifications do not affect the growth rate

Both differential interaction forces (25) and different growth rates (31) can induce cell sorting. We investigated whether deletion of the enzymes responsible for T4P post-translational modification affected the generation time of gonococci. Using a previously established assay for measuring the growth rate despite autoaggregation (16), we found that deletion of *pglO* or*pptA* and switching *pglE* into the ON-state did not the affect growth rate (Fig. 3).

**Fig. 3.**
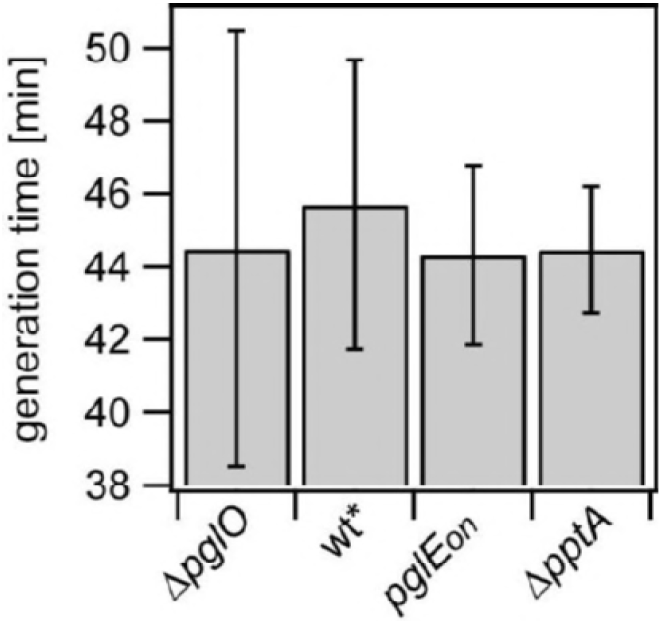
Pilin post-translational modification does not affect the generation time. The generation times of strains *mcherry-* expressing wt* (Ng170) and *gfp*-expressing *∆pglO* (Ng140), *pglE_on_* (Ng157), and *∆pptA* (Ng142) were determined during exponential growth on agar plates. Error bars: standard deviations form ≥ 90 colonies for each strain.

### The dynamics of shape relaxations is strongly affected by phosphoform and glycoform modifications

We found that the T4P-mediated *p_bound_* between bacteria and the rupture forces are affected by different pilin post-translational modifications. It was conceivable, therefore, that materials properties were affected. We characterized shape relaxations of microcolonies subsequent to fusion events (Fig. 4). To this end, gonococci were inoculated into a flow chamber. When oxygen was depleted, bacteria disassembled colonies (5). When oxygen was replenished by adding fresh medium, colonies re-formed within seconds. The oxygen-depletion step ensures that colonies form de-novo. When bacteria are freshly inoculated, then pre-existing colonies may contain a core in which bacteria are irreversibly bound (5), affecting the shape relaxation dynamics. After oxygen replishment, colony fusions were frequently observed. Shapes of wt* colonies relaxed considerably faster towards its spherical shape compared to the *pglE_on_* and *∆pptA* strains (Fig. 4a). To quantify relaxation dynamics, we analysed the relaxation from an ellipsoid shape to a spherical shape (Fig. 4a) assuming that the bacterial colonies behave like hard-sphere liquids, as described previously (17). In short, we assume that the energy change due to the reduction of surface area is balanced by the work of viscous deformation and use a fluid model of fusion dynamics to describe the shape changes during the fusion of two colonies (29). In this model, the initial shape of the fusing colony can be described by an oblate ellipsoid with the ratio of the minor axis *b* and major axis *a f* = *b/a* and its volume *v* = 4/3 *πab^2^*. Then the rate of deformation of the drop is

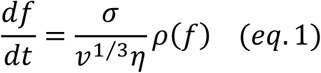

where *σ* is the surface tension and *η* the viscosity. *ρ*(*f*) is described in the Methods section. If at time *tc* an ellipsoid of cells has an axial ratio *f*_0_ and by time *t* it reaches *f* then

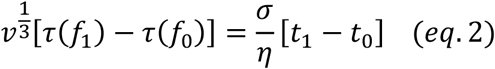

τcan be obtained by numerical integration of *ρ*(*f*) (Fig. 4b).

**Fig. 4.**
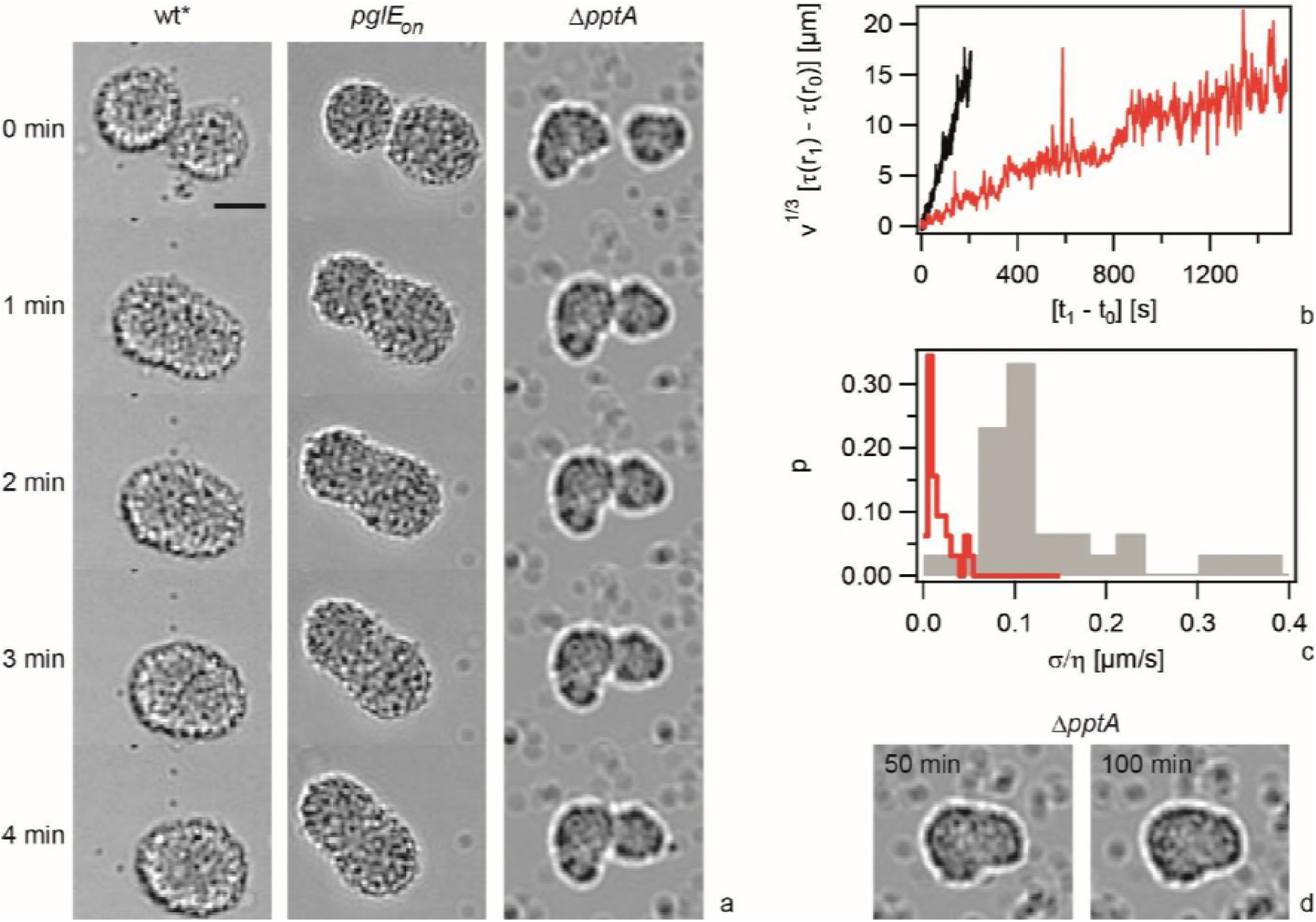
Post-translational modification of pilin strongly affects the dynamics of microcolony shape relaxations. a) Time-lapse of colony fusion events. Scale bar: 10 μm. b) *v*^1/3^[*τ*(*f*) – *τ*(*f*_0_ = 0.71)] as a function of time from fusion events shown in a). Black: wt* (Ng151), red: *pglE_on_* (Ng157). c) Distribution of ratio between surface tension *σ* and viscosity *η* obtained from eq. 2. Grey: wt* (Ng151), red: *pglE_on_* (Ng157)). N > 30 colonies for each condition. d) *∆pptA* (Ng142) fusion event after 50 min (~ tgen) and 100 min (~ 2 tgen).

For the wt* strain we obtained an average (± se) ratio between surface tension *σ* and viscosity η of (σ / η)_wt*_ = (0.15 ± 0.02) μm s^-1^ (Fig. 4c). The ratio is considerably smaller for the *pglE_on_* strain with (σ / η)_pglEon_ = (0.022 ± 0.005) μm s^-1^. Colonies formed by the *∆pptA* strain fused and the shapes of the colonies slowly relaxed towards spherical shapes (Fig. 4a). However, the time scale at which shape relaxations occured was slower than the generation time (Fig. 4d). Under these conditions, we cannot distinguish between effects caused by T4P-mediated re-organization of the colony and by duplication of bacteria; therefore, shape relaxations in the *∆pptA* strain were not further analysed.

We conclude that the presence of the diNAcBac-Gal-Gal trisaccharide slows down the colony relaxation dynamics, indicating that pilin glycoform status matters. The absence of the phosphoform modification at S^68^ led to an even more severe effect on colony relaxation dynamics; colonies did not relax towards a spherical shape within one generation.

## Discussion

### Different post-translational modifications of pilin affect the T4P-T4P mediated interaction forces between cells

T4P pilins are post-translationally modified in different bacterial species (32-35). We showed that deletion of genes encoding of enzymes responsible for adding phosphoform-modification at S^68^ as well as the glycoform modification at S^63^ (21) enhanced the rupture force between T4P emanating from different cells. Based on molecular dynamics simulations, it has been suggested previously for *N. meningiditis*, that inhibition of the phosphoform-modification is likely to enhance the contact area between T4P that are aligned side by side (24). This may explain the increase in rupture force observed here. In the strain used in our study, the pili bears a di-saccharide modification. We found that activation of the enzyme responsible for attaching a third saccharide, let to an increased rupture force. The molecular basis of this increase remains unclear.

### Change in T4P-mediated attractive force can explain the mesoscopic properties of microcolonies

The rupture forces and the probabilities that two bacteria are bound quantify the attractive force between two bacteria interacting via T4P-T4P binding. Thus, we can directly link attractive forces at the molecular level to collective behavior. Remarkably, the differences of rupture forces are fairly small between the wt and the mutants investigated in this study. When the rupture force between T4P increases, we expect that the probability that two cells are attached to each other via T4P increases. Indeed, the average probabilities that two bacteria are bound by T4P differ from the wt by ~ 45 *%* for the *∆pptA* and the *pglE_on_* strains (Fig. 2b). The differential strength of adhesion hypothesis (25) would predict that bacteria use T4P retraction to move with respect to each other and that bacteria preferentially move in the direction where rupture is least likely. Therefore, the sorting behavior observed in Fig. 1 can be readily explained; because the probabilities in the *∆pptA* and the *pglE_on_* strains are higher compared to the wt, the wt segregates to the surface of mixed colonies. Together with our previous study showing that inhibition of pilin glycosylation by deletion of *pglF* causes cell sorting (16), we can draw the general conclusion that inhibition of pilin glycosylation and phosphoform modification induces segregation of the wt cells to the surface of mixed colonies.

In addition to the robust sorting phenotype, we have shown that pilin post-translational modifications have severe effects on the dynamics of shape relaxation in gonococcal microcolonies. The relaxation time from an ellipsoid to a sphere after colony fusion is proportional to the ratio between surface tension and viscosity (29). Activation of PglE increased the relaxation time 10 fold. Still, shape relaxations from a dumbbell to a sphere occurred within on generation, suggesting liquid-like behavior. Inhibition of phosphoform-modifications showed an even more severe effect on the dynamics of shape relaxations. The relaxation time was larger than the generation time of ~ 45 min. As a consequence, most microcolonies did not assume a spherical shape. We expect that increasing surface tension enhances the relaxation rate whereas increasing viscosity lowers the relaxation rate. While we have not determined both quantities independently, we can estimate the surface tension knowing the rupture forces between T4P and the probabilities that bacteria are connected to each other by T4P. The experimental values are F_rupture wt*_ ≈ 51 pN, F_rupture pglEon_ ≈ 55 pN, and F_rupture pptA_ ≈ 64 pN. Please note the forces exceeding 80 pN were not measured accurately. The probabilities that bacteria are connected to each other by T4P p_bound wt*_ ≈ 0.15, p_bound pglE_on__ ≈ 0. 22, and p_bound pptA_ ≈ 0.24. These values enable us to estimate the surface tension *σ* = *F*∆*x* /∆*A* where *F∆x* is the work required to increase the surface area by *∆A*. The work required for moving one bacterium from the bulk to the surface by the distance *D* is estimated as follows. The diameter of a bacterium is D ≈ 0.7 μm, the average number of T4P per wt cell is N ≈ 7 (36), and the increase in surface area is ∆A ≈ ρ(D/2)^2^ ≈ 0.4 μm^2^. We assume that half of the pili are not bound when the bacterium resides at the surface. Together, we estimate

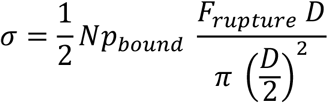

and obtain *σ*_wt*_ ≈ 5 · 10^-5^ Nm^-1^ σpglE_on_ ≈ 8 · 10^-5^ Nm^-1^ and σ_pptA_ ≈ 10^-4^ Nm^-1^. Using the experimentally determined values for *σ*/*η*, we find η_wt*_ ≈ 300 Nsm^-2^ and η_pglEon_ ≈ 3600 Nsm^-2^. This estimate suggests that increasing the attractive interaction between bacteria by altering the pilin posttranslational modification moderately enhances the surface tension and strongly enhances the viscosity. We note that the laser tweezers assay only samples anti-parallel binding between T4P, i.e. the tips of the pili are situated at the opposite ends of the complex. Within microcolonies, T4P of different cells may also assemble in parallel and the binding characteristics for this geometry are unkown. Most likely, the attractive interactions in the *∆pptA* strain were high enough to inhibit cellular rearrangements towards a spherical shape during one generation. As a consequence, the bacterial colonies do not exhibit liquid-like dynamics. Interestingly, meningococci generating non-glycosylated pilin form microcolonies with undefined shapes (22), consistent with strongly increased colony relaxation times. Taken together, different pilin post-translational modifications very strongly affect the materials properties of microcolonies formed by *Neisseria species*.

## Conclusion

We showed that different pilin post-translational modifications affect the pilus-mediated attractive interactions between bacteria. Quantitatively, these molecular effects are moderate. However, they have a strong impact on the materials properties of microcolonies. Interestingly, our laboratory strain showed the lowest attractive forces compared to all mutants investigated. As a consequence, wt* cells segregate to the surface of mixed microcolonies. In the laboratory environment, this behavior may be beneficial, because surface-dwelling bacteria have a selective advantage (18, 37). Moreover, shape-relaxations in fluid colonies occur fast. This behavior can be advantageous during colonization in confinement such as blood vessels (15). Different materials properties may be advantageous under different environmental conditions. T4P employ a plethora of mechanisms for controlling the primary structure of the pilin, the T4P density, and the status of post-translational modification by localized hypermutations (26). All of these mechanisms affect the attractive force between bacteria and enable them to switch their materials properties within hours.

## Author contributions

RZ, TC, NK, BM designed research; RZ, TC, NK, AW performed research; MK contributed bacterial strains; RZ, TC, NK, AW analyzed data; RZ, TC, AW, MK, BM wrote the manuscript.

## Acknowledgements

We thank the CECAD imaging facility of support with confocal microscopy, the Maier lab for helpful discussions, and the Deutsche Forschungsgemeinschaft for funding through grant MA3898.

